# Restoration of temperate forestry-drained fens brings back key species and an ecosystem function

**DOI:** 10.64898/2026.01.01.697300

**Authors:** Johannes Merz, Franziska Willems, Lena Lerbs, Anna Bucharova

## Abstract

1. Fen peatlands act as carbon sinks and are hotspots for specialised species, but a large proportion of fens have been drained for agricultural or forestry use. Drainage turns peatlands from carbon sink to carbon source, and negatively affects specialised biodiversity. Although restoration by rewetting can partially improve ecosystem functions, the successful recovery of specialised biota depends on previous land use. Rewetting agriculturally used fens often results in an ecosystem that does not resemble the original fen. The vegetation recovery of rewetted fens used for forestry is more promising, but our knowledge of vegetation development comes from boreal fens. It is unclear whether vegetation can successfully recover after the rewetting of forestry-drained fens in the temperate zone.
2. Our study focused on the recovery of vegetation and peat accumulation in small fens in temperate Europe, located in a large forest complex in Germany, that had been drained for forestry and restored by rewetting over the 38 years prior to the study. We compared the restored sites to degraded and near-natural ones. We recorded vegetation composition and measured peat depth.
3. We demonstrate that peat layers continuously increased with time since restoration, possibly indicating peat accumulation, which is crucial for converting drained fens back into carbon sinks. The vegetation composition of the restored sites became increasingly similar to that of the near-natural sites over time, with the steepest change occurring during the first 10 years after restoration. Nevertheless, even after decades, the vegetation did not reach the quality of near-natural sites.
4. *Synthesis and applications:* Our results highlight that rewetting forestry-drained temperate fens can lead to partial vegetation recovery and jump-start peat formation.

## Introduction

Habitat degradation is a pervasive phenomenon with negative consequences for humans and biodiversity (IPBES, 2018). Biodiversity maintains functional ecosystems and their services we all depend on (Hong et al., 2022). Ecosystem restoration is a major strategy to accelerate the recovery of ecosystems, their biodiversity, and the services they provide (Atkinson et al., 2022, Leclère et al., 2020).

An important ecosystem that has been severely degraded by human activities is fen peatland (Tanneberger et al., 2021). Fen peatlands are wetland ecosystems fed by groundwater and dominated by highly specialized peat mosses and sedges. The biomass of these plants forms peat, that is litter accumulated in waterlogged conditions, where it is prevented from oxygen access and consequent decomposition (Joosten et al., 2017, Joosten & Clarke, 2002). Fens are thus substantial carbon sinks (Leifeld & Menichetti, 2018). Fen peatlands also retain water in the peat and act as a buffer against extreme weather events (Ahmad et al., 2020, Gatis et al., 2023). However, natural waterlogged fens are not suitable for intensive forestry or agriculture. To make these areas profitable, humans have drained them. While drainage allowed planting trees or growing crops, it completely altered the fen peatland ecosystem and led to profound degradation (Tanneberger et al., 2021, Joosten & Clarke, 2002).

Degraded fen peatlands cannot support unique fen ecosystem services and have lost most of their specialized species (Andersen et al., 2017, Lamers et al., 2015, Joosten et al., 2017). Exposing formerly waterlogged peat to aerial oxygen leads to decomposition of organic matter and loss of peat volume, which results in extreme levels of greenhouse gas emissions (Freeman et al., 2001, Mrotzek et al., 2020). Mineralisation of nutrients in the degraded peat can cause internal eutrophication (Smolders et al., 2006). These changes often permanently alter the characteristics of the ecosystem to which fen specialized biota is adapted (Lamers et al., 2015, Klimkowska et al., 2019). As a result, the biodiversity of fens is lost (Klimkowska et al., 2010, Lamers et al., 2015).

Fen restoration typically consists of rewetting, with the aim to restore wet and nutrient-limited conditions and to support the establishment of typical fen vegetation (Joosten 2021, Allan et al., 2024, Tanneberger et al., 2021). Rewetting can immediately stop greenhouse gas emissions and may lead to new carbon sequestration through peat accumulation (Mrotzek et al., 2020, Günther et al., 2020). Although rewetting can cause methane emission, the long-term climate balance of rewetting is always positive (Günther et al. 2020). Restored fens can also buffer extreme rainfall better than degraded fens (Ahmad et al., 2020, Gatis et al., 2023). However, the trajectory of vegetation recovery after rewetting depends on land use after drainage (Lamers et al., 2015). Fen peatlands in temperate Europe have been mainly drained for agricultural use (Joosten et al., 2017, Tanneberger et al., 2021). Here, rewetting often mobilises high levels of phosphorus in the degraded peat (Zak et al., 2007, Smolders et al., 2006), which supports growth of tall marsh plants (helophytes) that prevent establishment of less competitive fen specialists (Kreyling et al., 2021, Roth et al., 1999). With that, the helophytes may trap succession in alternative states which can result in novel ecosystems that do not resemble original fens (Roth et al., 1999, Kreyling et al., 2021). In boreal zone, fens have been mainly drained for forestry. Here, the peat degradation was not as severe as in former fens used for agriculture, and rewetting can return the geochemistry and vegetation characteristics on the recovery trajectory towards conditions at natural sites (Elo et al., 2024). In any case, restored fens only rarely reach vegetation composition of undegraded fens, and differences can be detected even after decades (Kreyling et al., 2021, Andersen et al., 2017, Elo et al., 2024).

Much of our knowledge on the vegetation recovery in restored fen peatlands in Central Europe comes from large agricultural drained fen complexes (e.g. Kreyling et al., 2021). However, also in Central Europe, there are smaller fens that have been drained for forestry, and are now restored by rewetting (Tanneberger et al., 2021, Andersen et al., 2017). It is unclear if forestry-drained fens in Central Europe follow the same recovery pathway as forestry-drained boreal fens, particularly because substantial differences in climate, more intensive use of the landscape, and higher level of nitrogen through atmospheric deposition in Central Europe (EEA, 2024, EEA, 2015). Further, small fen ecosystems may differ from their larger counterparts because ecosystem functioning and biodiversity relationships can change non-linearly with size (Gonzalez et al., 2009). Ecosystem size significantly affects the recovery after restoration, and we do not know whether we can directly transfer knowledge from large fen restoration to restoration of small fens (Moreno-Mateos et al., 2012).

To address this research gap, we focused on restoration of small, forestry-drained fens located in a large forest complex in Central Germany, Europe. We recorded vegetation composition, including mosses, and measured peat depth as a proxy of peat formation at degraded, near-natural and restored fens. The time since restoration varied between 2 and 38 years, which allowed us to employ space-for-time approach. We hypothesize that (1) restored fens will host more fen-specialized species, and support more fen-like ecosystem functions (specifically, peat formation) than unrestored sites; (2) restored fens will become increasingly similar to near-natural fens over the course of nearly 40 years since restoration; and (3) despite this trend, even the oldest restored fens will differ from the near-natural fens in all studied aspects.

## Methods

### Site description

All fens are located within a large forest complex (Burgwald) in Hesse, Germany (**Figure 1**). The fens in the Burgwald are nutrient-poor sloping fens with a flow of groundwater (percolation) (Joosten et al., 2017, Küchler, 2017, Küchler, 2018). Historically, they were used as meadows, but most of them were drained and afforested with spruce in the first half of the 20th century. Since 1985, the local administration has been continuously restoring drained fens by clear-cut and rewetting. In our study, we included all restored fens in the area that were classified as sloping fens and for which we could obtain information on the year of restoration. Specifically, we included 20 fens that had been restored over 2-38 years prior to the study and thus varied in successional state. We compared the restored sites with four near-natural sites that had not been successfully drained and afforested, and two unrestored sites. The number of near-natural and unrestored fens is rather small, but these were all fens of this type that were available in the area. Each individual fen is referred to as a “site” in further text, in total, our study comprised 26 sites. We treated sites as independent if they differed in year of restoration and/or if they were spatially separated by roads, dams, or forest.

**Figure 1:**
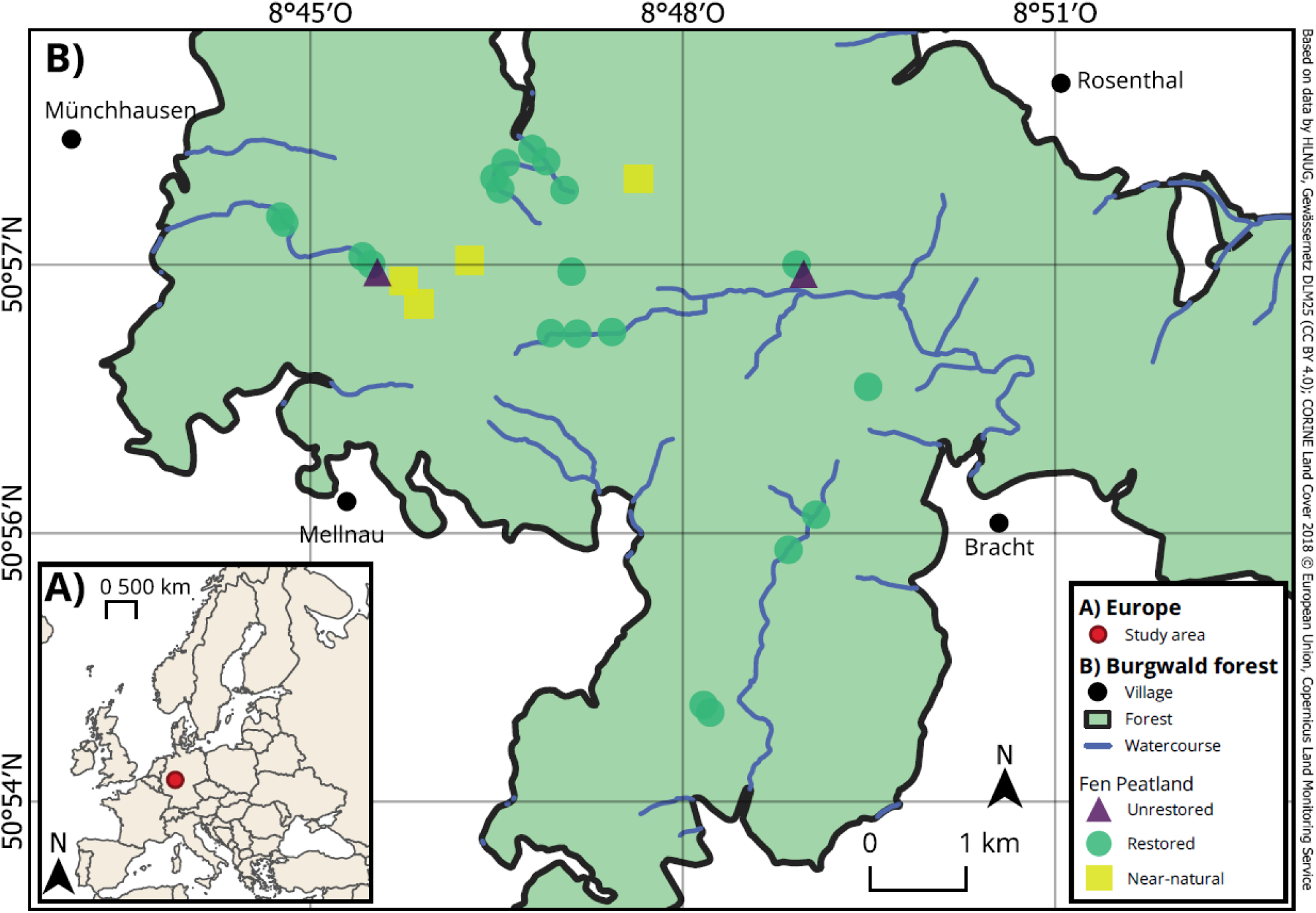
A) Location of study area within Europe. B) Map of the study sites in the Burgwald forest, Hesse, Germany

### Data collection

At each site, we randomly placed three transects, perpendicular to the water flow. Each transect was centred on the valley-axis of the fen, was 9m long and consisted of five 1m² (1x1 m) plots with 1m between plots. In each plot, we identified all vascular plants, bryophytes, liverworts and macro fungi, and recorded their cover in percentage. We identified all taxa to species level, except *Rubus spec, Callitriche spec., Hypericum spec.* which we identified to genus level. To get an approximate peat depth (cm), we pushed a rod into the centre of each plot until it hit the subsoil. We sampled the vegetation from mid-May to beginning of June 2024. We obtained the year of restoration from the local forestry office. For 9 restored sites, the exact year of restoration was not known. We thus estimated the year of restoration based on aerial photographs taken between 1975 and 2020 in on average 4-year intervals, as provided by the Hessian State Office for Land Management and Geoinformation.

### Statistical Analysis

Prior the analysis, we aggregated plots within a transect because transects, rather than individual plots, is a representative sample of the site. To do this, we calculated the mean cover for each individual species, and the mean peat depth across the five plots of the transect. In further analysis, one replicate is a transect. To quantify the cover of fen specialist species, we summed the cover of fen specialist species per transect. As fen specialist species, we considered species that are diagnostic for the habitat type ‘poor fens’ according to the European Nature Information System (category Q22). That means they are typical and indicative of this habitat type across Europe (Chytrý et al., 2020, Chytrý et al., 2024). See **Table S1** for a list of all fen specialist species that occurred in our study.

In the first step, we assessed differences in plant community composition between degraded, near-natural and restored sites. To capture differences along the successional gradient, we divided the restored sites to young ones (younger than 10 years) and old ones (older than 20 years). This covered all restored sites, because no restored site in our study was between 10 and 20 years old. We thus had four types of sites: degraded, young restored, old restored and near-natural. To assess differences in plant community composition among these site types, we used multivariate analyses using the vegan package in R (version 2.6.10, Oksanen et al., 2025). Compositional dissimilarity between sites was quantified using the Bray-Curtis distance metric. We then performed non-metric multidimensional scaling (NMDS, metaMDS function) with two dimensions to visualize community composition patterns across site types. Ordination quality was assessed using stress values, with values <0.2 indicating acceptable representation of the data. Subsequently, we tested for significant differences in community composition among site types using permutational multivariate analysis of variance (PERMANOVA; adonis2 function, *pairwiseAdonis* package version 0.4.1, Martinez Arbizu, 2020) with 999 permutations. To account for the nested study design (transects within sites), we aggregated community composition data to the site level by calculating mean species cover per site prior to PERMANOVA analysis. This approach avoids pseudoreplication arising from treating transects as independent observations.

To identify which site types differed from one another, we subsequently conducted pairwise PERMANOVA comparisons also using the *pairwiseAdonis* package. We assessed the assumption of homogeneity of multivariate dispersions using the betadisper function (*vegan* package), which calculates the distance of each sample to its group centroid. We tested differences in dispersion among groups using permutation tests (see supporting information **Figure S1**).

In the second step, we tested whether 1) peat depth, 2) total species richness, 3) richness of fen specialist species and 4) cover of fen specialist species differed between restored, unrestored and near-natural sites. Note that the age of restoration was implemented in the next step as a continuous variable (see below). We used generalized linear mixed effect models (glmmTMB function, *glmmTMB* package version 1.1.12, Brooks et al. 2025) with site type (unrestored, restored or near-natural) as fixed effects. We included a random intercept to account for clustering or non-independence of measurements of transects within the same site. All models shared the following core structure:

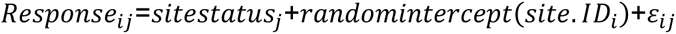

Full notation of the four final models is given in supporting information **S1**. To meet the assumptions of normality and homoscedasticity, response variables were checked for appropriate distributions: For the peat depth and specialist cover models, the response variables were log(x+1) transformed prior to analysis to improve linearity, stabilize variance and avoid predictions of negative values. For these two models, variance was notably lower in restored sites. To account for this heteroscedasticity, we included a variance function via the dispformula argument in glmmTMB, allowing residual variance to be estimated separately for unrestored, restored, and near-natural sites. This approach ensured robust standard errors and inference for the fixed effect. For the peat depth model, some differences in variance between the groups remained (see supporting information **Figure S2**). We modelled the specialist cover values with a beta distribution, using proportions (continuous data bounded between 0 and 1). For easier interpretation, values were expressed as percent cover in the result tables and figures. The fixed effect of site status was tested for significance using a Type II Analysis of Variance (package *car,* function Anova, Fox et al. 2024). Subsequently, we used a Tukey test for pairwise comparison between unrestored, near-natural and restored sites (package *emmeans* version 1.11.2, function emmeans, Lenth 2025). The explanatory power of all GLMMs was quantified using the marginal (R^2^m) and conditional (R^2^c) R^2^ values (package *performance*, version 0.15.2, function r2_nakagawa, Lüdecke et al., 2025). For the heteroscedastic model analyzing peat depth, R^2^ was estimated from a corresponding homoscedastic model (dispformula=∼1) that retained the same mean (fixed and random) structure, as the standard R^2^ methods assume a single residual variance. For model diagnostic plots see supporting information, **Figures S2- S5**.

In the third step, we tested whether 1) peat depth, 2) total species richness, 3) richness of fen specialist species and 4) cover of fen specialist species in the restored transects changed with time since restoration. To analyze how total species richness and fen specialist richness, as well as peat depth, changed over time, we used Generalized Additive Models (GAMs) (package *mgcv*, version 1.9.3, gam function, Wood 2025). We used GAMs to allow for nonlinear changes over time. Model evaluation showed that these nonlinear smooth (s) effects were the best fit for species richness, fen specialist richness, and specialist cover proportion, but a linear trend was retained for peat depth. To analyze the specialist cover proportion, we employed a Shape-Constrained Additive Model (SCAM), using a monotonically increasing (’mpi’) smooth for year (package *scam*, version 1.2.19, function scam, Pya 2025). We applied this constraint based on the ecological assumption that, after restoration, specialist cover is reasonably expected to remain the same or increase over time. Peat depth was modeled with a Gaussian distribution, species richness and fen specialist richness used a Poisson distribution, and specialist cover proportion employed a quasibinomial distribution. All models controlled for site-to-site differences by including site ID as a random effect and shared the following core structure:

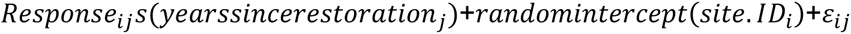

Full notation of the four final models is given in supporting information **S2**.

We quantified explained variance for each model by reporting fixed-effects and total model fit. We estimated “fixed-only” R^2^ by refitting each model without random-effect smooths and extracting the adjusted R² from the refit. “Total” fit for GAMs was taken as the adjusted R² from the original model including random-effect smooths. To calculate the R^2^ values we used the r2_nakagawa function in the *performance* package (Lüdecke et al., 2025). To assess whether model residuals were spatially autocorrelated, we tested for spatial autocorrelation using Moran’s I for all four GLMMs. Spatial weights were constructed using 12 nearest neighbours (package *spdep*, version 1.4.2, function knearneigh, Bivand & Wong 2018), based on UTM Zone 32N projected coordinates. No significant spatial autocorrelation was detected in any of the four models (all p > 0.05), suggesting that the random effect structure adequately accounted for the spatial structure of the data.

All analyses were performed in the R statistical environment (R Core Team 2024). We used the package *ggplot2* (version 3.5.2) for data visualization and the package *patchwork* (version 1.3.0) for arranging the plots (Wickham, 2016, Pedersen, 2024). In all models, we graphically evaluated whether model assumptions are met (Zuur et al., 2010). For evaluating GAMs we used the *gratia* package (version, 0.11.1, Simpson 2024) for the other models we used the *DHARMa* package (version 0.4.7, Hartig, 2024). For the resulting model diagnostic plots see supporting information **Figures S6 - S9**.

## Results

Non-metric multidimensional scaling (NMDS) of transect-level community composition based on Bray–Curtis dissimilarities produced an acceptable two-dimensional solution (stress = 0.172), indicating a reliable depiction of differences among transects. The ordination revealed a clear compositional gradient from unrestored to restored sites, progressing toward near-natural reference conditions (**Figure 2**). Notably, older restored sites clustered closer to near-natural communities than younger restored sites, suggesting that restoration promotes gradual recovery of species composition over time and that ecological trajectories of restored habitats move progressively toward conditions at near-natural sites. Community composition differed significantly between site types (PERMANOVA: R² = 0.41, F = 4.995, p < 0.001), explaining approximately 40% of the multivariate variance. All pairwise contrasts were significant except that between unrestored and near-natural sites (999 permutations, see **Table 1**), likely reflecting low statistical power due to small sample sizes rather than ecological similarity, as these groups showed the strongest separation in ordination space.

**Figure 2:**
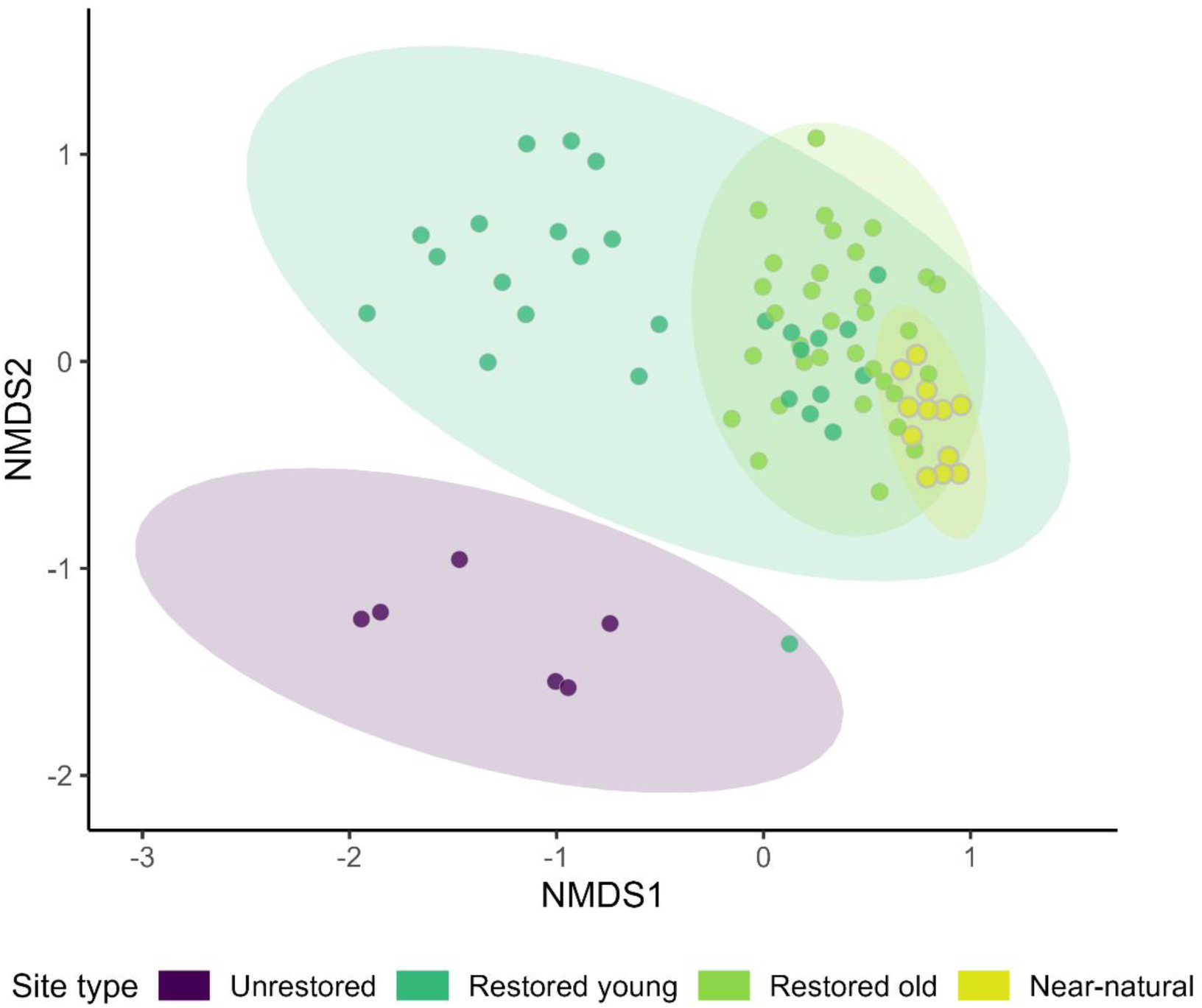
Non-metric multidimensional scaling (NMDS) of community composition based on Bray–Curtis dissimilarities. Points are individual transects, colored by site type; shaded polygons show 95% normal-data ellipses for each group. Stress = 0.172, indicating a good low-dimensional representation.

**Table 1:** Pairwise PERMANOVA comparisons of plant community composition among site types. R² indicates the proportion of variance explained by the contrast, F is the pseudo-F statistic, and p-values are based on 999 permutations of Bray–Curtis dissimilarities. Restored sites were differentiated between young (restoration less than 10 years ago) and old (restored more than 20 years ago). Significant differences are highlighted in bold.

| Group 1 | Group 2 | $R^2$ | $F$ | $p$-value |
| --- | --- | --- | --- | --- |
| Unrestored | Restored young | **0.25** | **3.03** | **0.019** |
| Unrestored | Restored old | **0.38** | **6.74** | **0.017** |
| Unrestored | Near-natural | 0.66 | 7.6 | 0.067 |
| Restored young | Restored old | **0.21** | **4.82** | **0.004** |
| Restored young | Near-natural | **0.33** | **5.41** | **0.009** |
| Restored old | Near-natural | **0.27** | **4.78** | **0.002** |

Vegetation composition was significantly more variable within the restored and unrestored groups than within the near-natural reference sites (Multivariate test of dispersion ANOVA: F = 12.361, p < 0.001) (**Figure S1**), though this result should be interpreted with caution given the unequal number of sites per group. This difference means the PERMANOVA results reflect not only a shift in the average composition but also a difference in the consistency of the vegetation at these sites.

Fen restoration significantly influenced key ecosystem function and vegetation parameters. At restored sites, peat depth as well as the richness and cover of fen specialist species were higher than at unrestored sites (Table 2, Figure 3). Although the number of specialist species in restored sites approached those in near-natural fens, their cover remained lower. Total species richness was slightly higher in restored than in near-natural sites(Table 2, Figure 3). Restoration status (unrestored, restored, or near-natural) explained between 16% - 67 % of the variance in the models (marginal R²), while including random site effects (conditional R²) increased the explained variance to 53 – 99 %. This suggests that although restoration status was an important predictor, a substantial proportion of variation in peat depth and vegetation characteristics was due to inherent differences among sites. However, Moran’s I indicated no significant spatial autocorrelation in the residuals of any of the four models (all p > 0.05), confirming that the results are not confounded by spatial structure.

**Figure 3.**
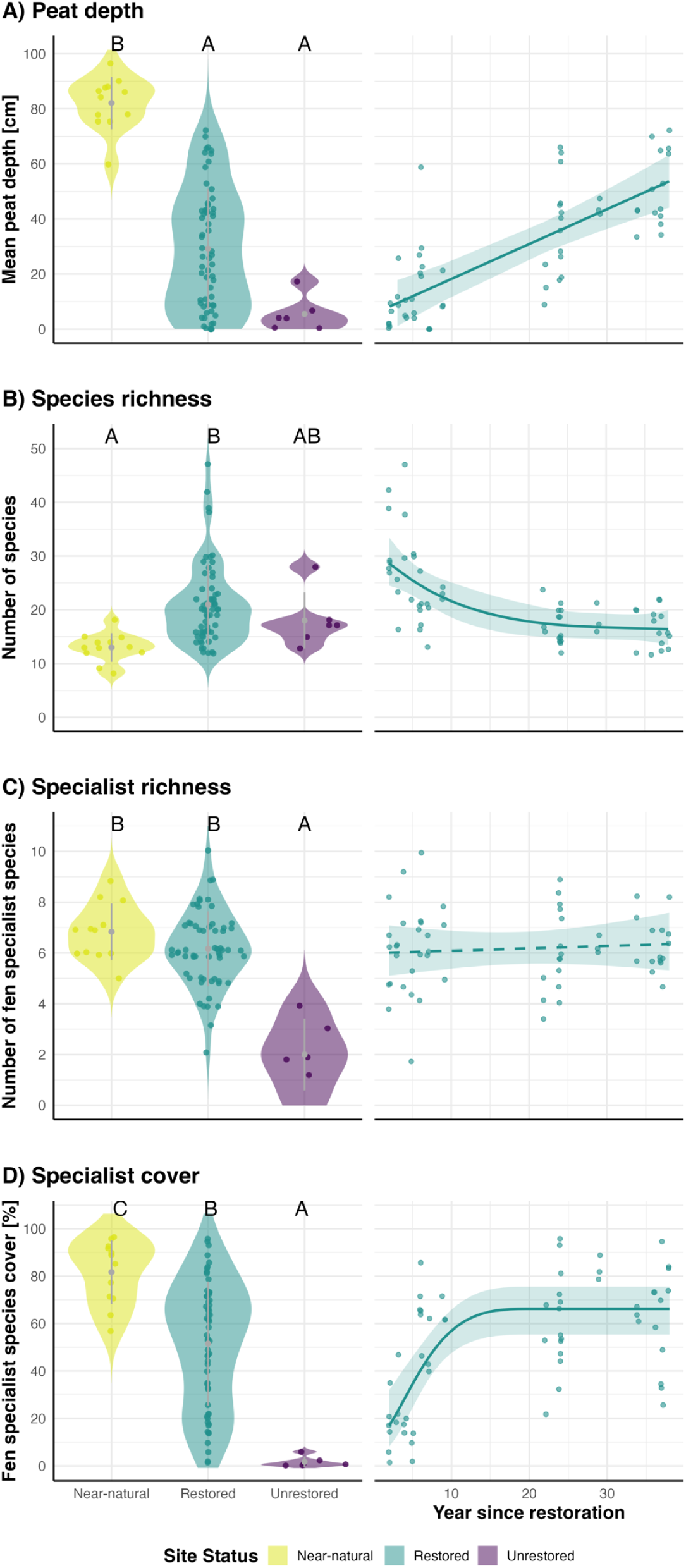
Peat depth (A), species richness (B), fen specialist species richness (C), and fen specialist species cover (D) at unrestored, near-natural and degraded sites, and development of the same variables across time since restoration at restored plots. **Left:** Violin plots showing the distribution of each variable in near-natural, restored, and unrestored sites. Grey dots and whiskers represent mean ± SE. Letters above violins indicate significant differences between site types (A, B; Tukey post-hoc test, α = 0.05). **Right:** Scatterplots of the mean values in restored sites over years since restoration, with fitted trend lines and 95% confidence intervals. Individual points represent transect-level measurements.

**Table 2:** Summary of Linear Mixed-Effects Model (LMM) results testing the effect of site status on peat depth and vegetation characteristics. Fixed Effect Test reports the overall significance of site status (unrestored, restored or near-natural), based on an ANOVA using a χ² Tests. P-values less than 0.05 are shown in **bold.** Estimated marginal means (EMMs) not sharing the same lowercase letter (a, b) within a row are significantly different (p<0.05) based on Tukey-test post-hoc comparisons. CI indicate 95% confidence intervals of the EMMs. R^2^m (Marginal R^2^) represents the variance explained by the fixed effect (site status), and R^2^c (Conditional R^2^) represents the variance explained by both fixed and random effects.

| Response variable | Fixed Effect Test |  | Estimated Marginal Means (95% CI) |  |  | Model Fit |  |
| --- | --- | --- | --- | --- | --- | --- | --- |
| | $\chi^2$ | *P-value* | *Unrestored* | *Restored* | *Near-natural* | $R^2m$ | $R^2c$ |
| Peat depth [cm] | 11.91 | **0.003** | 3 (0 – 27) a | 18 (10 – 33) a | 82 (23 – 279) b | 0.30 | 0.92 |
| Species richness [No.] | 6.42 | **0.040** | 18 (9 – 28) ab | 21 (18 – 24) b | 13 (6 – 20) a | 0.16 | 0.69 |
| Specialist richness [No.] | 38.56 | **0.001** | 2 (1 – 3) a | 6 (6 – 7) b | 7 (6 – 8) b | 0.41 | 0.53 |
| Specialist cover [%] | 37.49 | **0.001** | 1 (0 – 7) a | 51 (40 – 62) b | 84 (66 – 93) c | 0.67 | 0.99 |

The effects of restoration changed with time since the measures were implemented. Peat depth showed a sustained and significant long-term increase, while vegetation composition mostly changed only during the first decade after restoration. (**Table S2, Figure 3A**). In contrast, total species richness declined initially – from almost 30 species on recently restored sites to a plateau of approximately 16–17 species on sites restored more than 20 years ago (**Table S2, Figure 3B**). Although the number of fen specialist species did not change with time since restoration (**Table S2, Figure 3C**), their cover increased steeply during the first decade after restoration and stabilized around 66 % (**Table S2, Figure 3D**). Variation among sites remained high (**Figure 3D**). Across most variables, the models successfully explained a large fraction of the observed changes, generally ranging from approximately 50% to 70% of the total variance. A notable exception was fen specialist richness, where the time since restoration had almost no consistent effect (fixed R²=−0.004). Temporal model results are summarized in **Table S2** in the supporting information.

## Discussion

Drainage is the main cause of fen peatland degradation and the loss of their specialized biodiversity (Joosten et al., 2017). Consequently, rewetting is the primary restoration strategy. Here we demonstrate that rewetting can rapidly restore vegetation and trigger peat accumulation in small, forestry-drained fens located within a large forest complex in Central Europe. While degraded sites hosted only very few fen-specialized species and these had low cover, restoration increased the number of specialists to levels comparable with those in near-natural sites. Their cover also gradually increased over time since restoration. Consequently, the vegetation composition of the restored sites became increasingly similar to that of the near-natural sites over time, but did not reach the same quality, even after decades.

The post-restoration development of vegetation towards the reference ecosystem contrasts sharply with the results reported from restoration of large Central European fens that were mainly drained for agricultural use. In these cases, vegetation of rewetted fens turned to locally novel ecosystem types dominated by tall vascular plants (helophytes), a habitat that does not resemble the original fen peatland (Kreyling et al., 2021). Agricultural use commonly leaves a legacy of soil compaction and eutrophication (Lamers et al., 2015), severely degrading the peat soil and hindering the development of nutrient-poor fens. Peatlands that were previously used for agriculture are also more likely to be embedded in agricultural landscapes and may be affected by ongoing nutrient input from the surrounding area (Hurkuck et al., 2014). In our study, we focused on small fens that had been used for forestry. Although forestry-drained fens may have been fertilized prior to tree planting (Haapalehto et al., 2017), the nutrient input was certainly lower than in agricultural use. Furthermore, the fens are surrounded by a large forest complex which limits nutrient input from adjacent land use. Previous studies on forestry-drained fens in Northern Europe have shown that rewetting can initiate succession towards the original ecosystem (Elo et al., 2024). In further discussion, we thus compare our results predominantly with studies from forestry-drained fens.

The community composition of restored fens became increasingly similar to that of near-natural fens over time. This trend was driven by an increase in the number and coverage of fen specialist species, primarily sedges (*Carex* spp.) and peat mosses (*Sphagnum* spp.). These species outcompeted the generalist species that had initially colonised the restored sites following disturbance. As the generalist species were lost, the total number of species decreased to a level comparable with that of near-natural sites. The loss of generalist species as succession proceeds is common in rewetted, forestry-drained fens (Haapalehto et al., 2017). Nevertheless, even nearly 40 years after restoration, the community composition of rewetted fens did not fully reach the quality of near-natural fens, which confirms the common paradigm in restoration ecology that restored ecosystems rarely reach the quality of pristine ones (Moreno-Mateos et al., 2012, Atkinson et al., 2022).

Although drained fens hosted only very few species typical of near-natural fens in the area, we observed all of these specialists at least at some restored sites. Such full recovery of fen-specialized flora is uncommon, even at restored forestry-drained sites (Haapalehto et al., 2017, Elo et al., 2024). One possible reason behind this is effective dispersal. The small size of the individual sites allowed for restoration of multiple sites (Bentley et al., 2022), which possibly recovered the connection between fen fragments and facilitated seed dispersal, a factor which is often limiting in fen restoration (Renton et al., 2012, Klimkowska et al., 2010). Without effective dispersal, such full return of fen specialists is unlikely as seed banks are often degraded after decades of drainage (Lamers et al., 2015, Klimkowska et al., 2010).

The vegetation recovery was particularly rapid during the first ten years after restoration, with only little change in the following decades. It is possible that the first ten years were sufficient to stabilize the community. Previous restoration studies of forestry-drained fens have mostly covered a maximum of ten years after rewetting and have shown rapid changes during this time (e.g. Haapalehto et al., 2017, Elo et al., 2024). The long-term dynamics of forestry-drained fens after rewetting remain largely unexplored. Studies on the rewetting of mainly large, agriculture-drained fens have shown that the vegetation remains largely unchanged for over 30 years (Kreyling et al., 2021).

The peat layer continuously increased with the age of restored sites. This might indicate new peat accumulation after restoration, as rewetting can jump-start peat formation even if a near-natural vegetation composition is not achieved (Günther et al., 2020, Kareksela et al., 2015). However, the total peat depth is also influenced by the pre-restoration peat depth. It is possible that sites that were restored more recently were more degraded prior restoration and had thinned layer of peat. Despite this cofounding factor, part of the peat depth is likely due to new peat accumulation because of the simultaneous return of peat forming specialist species (e.g Sphagnum). Peat formation restores the fens crucial function of fens as a carbon sinks (Günther et al., 2020), and new peat layers and a higher peat moss cover can improve the water retention (Ahmad et al., 2020, van de Koot et al., 2024).

An important limitation of our study is that we have no data on pre-restoration state. It is possible that selection of sites for restoration was not random, and the least degraded sites were restored first. This could affect our conclusions on peat depth, as discussed above. However, it is unlikely to affect our conclusion on temporal development of vegetation, because we observe continuous changes in specialist cover only during the first 10 years, followed by stagnation during subsequent 30 years. Possible better pre-restoration state of the oldest restored sites cannot explain this pattern.

In summary, we have shown that rewetting small, forestry-drained fens located in a large forest complex can be surprisingly successful. The restoration brought back target fen species and possibly initiated peat accumulation. Although the vegetation composition did not reach the quality at near-natural fens, the community was more similar to the reference ecosystem than we have expected based on literature from other restored fens in Central Europe. The vegetation development resembled the post-restoration changes of forestry-drained fens in boreal zone (Elo et al., 2024), probably because the absence of nutrient input from the surrounding landscape and less severe degradation in comparison to agriculturally drained fens. The small size of the fens and their location in valleys may have allowed their connection to the near-natural fens in the region, facilitating the dispersal of the target plant species. Our results highlight that small-scale restoration of forestry-drained fen peatlands is a promising strategy, especially given that small projects are often cheaper and easier to implement (Maynard, 2013).

## Acknowledgement.

We thank the Forestry Office Burgwald for providing documentation on the restoration projects, the Regierungspräsidium Gießen for issuing permits for field work in nature protection areas, and the AG Rettet-den-Burgwald for assistance.

## Notes

### Competing Interest Statement

The authors have declared no competing interest.

### Summary of Updates

Wording and literature edited in the whole manuscript. Method section updated, in particular permanova to account for non-independency of the plots. Figure 1 edited for clarity. One author added (Lena Lerbs)

