## supporting for "Restoration of temperate forestry-drained fens brings back key species and an ecosystem function"

### Supporting information

**Table S1.** List of all fen specialist species recorded at our study sites. Note that some taxonomic discrepancies exist between our dataset and the European Nature Information System (EUNIS; Chytrý et al. 2020, Chytrý et al. 2024). For the calculation of richness and cover of EUNIS Q22 diagnostic species, we used the species aggregates defined in EUNIS. This applies only to *Sphagnum recurvum* aggr. (which, according to the World Flora Online Consortium, comprises *S. angustifolium* s. str., *S. fallax* s. str., and *S. flexuosum* s. str.), *Sphagnum palustre* aggr. (represented only by *Sphagnum palustre* s. str.) and Molinia caerulea aggr. (represented only by *Molinia caerulea* s. str.). For all other taxa, our vegetation taxonomy follows the accepted names of the World Flora Online Consortium (2024).

| EUNIS Diagnostic species Q22 ´poor fen´ present at our study sites |
|---|
| *Agrostis canina* |
| *Aulacomnium palustre* |
| *Carex canescens* |
| *Carex echinata* |
| *Carex nigra* |
| *Carex nigra* |
| *Drosera rotundifolia* |
| *Eriophorum angustifolium* |
| *Molinia caerulea* s. str. |
| *Polytrichum commune* |
| *Sphagnum palustre* s. str. |
| *Sphagnum papillosum* |
| *S. angustifolium* s. str., *S. fallax* s. str., *S. flexuosum* s. str. |
| *Vaccinium oxycoccos* |
| *Viola palustris* |

**Multivariate analysis:**

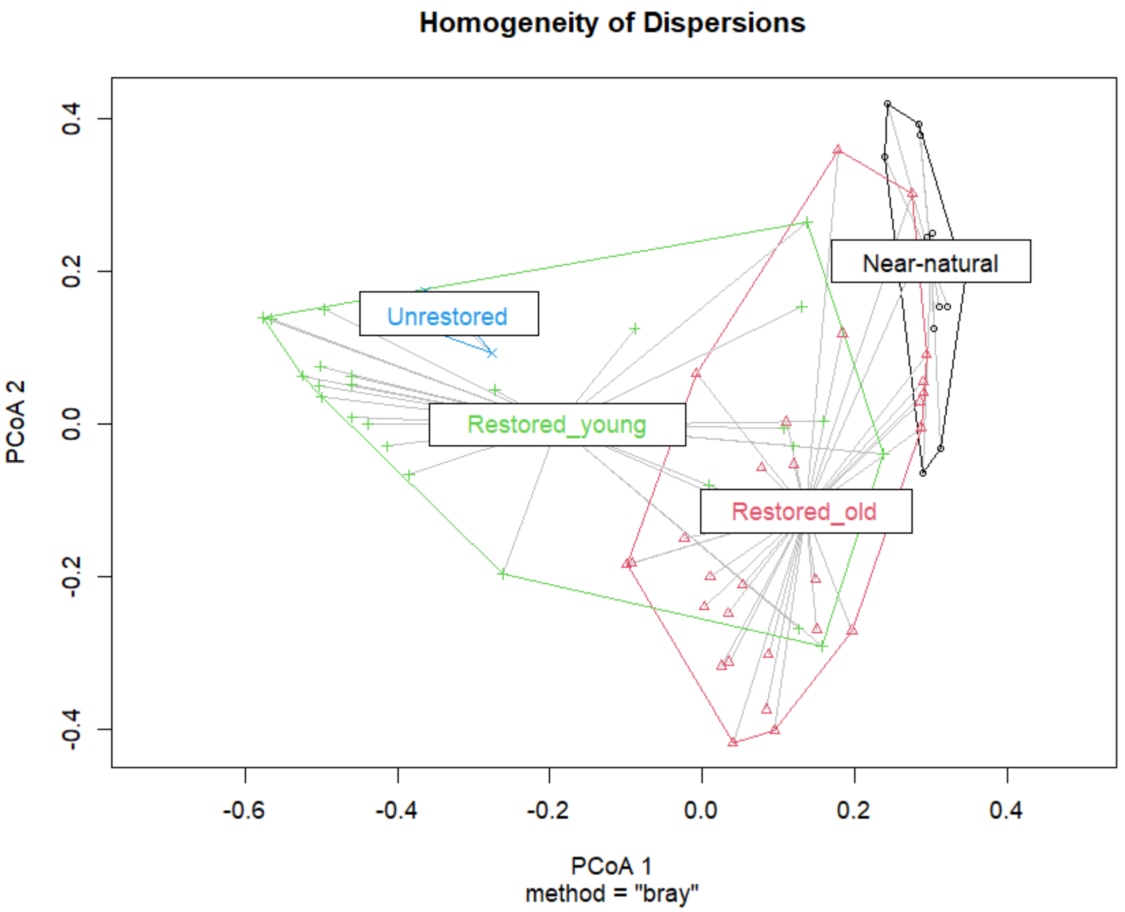


**Figure S1**: Tests of multivariate dispersion (betadisper) revealed significant differences in within-group variability among site types (ANOVA: F = 21.697, p < 0.001). Near-natural sites were less dispersed than both Restored sites and Unrestored **sites** (e.g., Near-natural − Restored_young: p < 0.001; Near-natural − Restored_old: p < 0.001), and Restored_old were less dispersed than Restored_young (p < 0.001), consistent with increasing community homogeneity through restoration. Points represent individual transects.

**Supporting information S1: Site Status Model notation**

Full model notation of model investigating the differences between unrestored, restored, and near-natural sites (called Site Status below). The four models were fitted using Generalized Linear Mixed Models (GLMMs) via the glmmTMB package (Brooks et al. 2025, McGillycuddy et al. 2025).

**1) Peat Depth Model**

The peat depth was log-transformed and modeled using a **Gaussian** distribution with a fixed effect of Site Status and a random intercept for Site ID. The residual variance ($\sigma^2$) was modeled as heteroscedastic, allowing it to vary by Site Status.

$$log(\text{peat depth}_i + 1) \sim N(\mu_i, \sigma^2_{\text{SiteStatus}})$$

$$\mu_i = \beta_0 + \beta_1 \cdot \text{SiteStatus}_i + u_{j[i]}$$

$$log(\sigma^2_{\text{SiteStatus}}) = \gamma_0 + \gamma_1 \cdot \text{SiteStatus}_i$$

Where $\mu_i$ is the predicted mean, $u_{j[i]}$ is the random intercept for Site ID j, and $\sigma^2_{\text{SiteStatus}}$ is the residual variance (dispersion parameter) specific to the level of Site Status. In all models $u_j$ was assumed to be normally distributed: $u_j \sim N(0, \tau^2)$, with $\tau^2$ being the random effect variance.

**2) Species Richness Model**

The total number of plant species was modeled using a **Gaussian** distribution ($\text{Species Richness}_i \sim N(\mu_i, \sigma^2)$) with a fixed effect of Site Status and a homoscedastic random intercept for Site ID: $mu_i = \beta_0 + \beta_1 \cdot \text{SiteStatus}_i + u_{j[i]}$

**3) Fen Specialist Species Richness Model**

The number of fen specialist species was modeled using a **Gaussian** distribution ( $\text{Specialist Richness}_i \sim N(\mu_i, \sigma^2)$) with a fixed effect of Site Status and a homoscedastic random intercept for Site ID: $mu_i = \beta_0 + \beta_1 \cdot \text{SiteStatus}_i + u_{j[i]}$

**4) Fen Specialist Species Cover Model**

The specialist cover proportion was modeled using a **Beta** distribution with a logit link with a fixed effect of Site Status and a random intercept for Site ID. The precision parameter ($\phi$) was modeled as heteroscedastic, allowing it to vary by Site Status.

$$\text{Specialist Cover}_i \sim \text{Beta}(\mu_i, \phi_{\text{SiteStatus}})$$

$$logit(\mu_i) = \beta_0 + \beta_1 \cdot \text{SiteStatus}_i + u_{j[i]}$$

$$log(\phi_{\text{SiteStatus}}) = \gamma_0 + \gamma_1 \cdot \text{SiteStatus}_i$$

Where $\mu_i$ is the predicted mean proportion on the probability scale, logit($\mu_i$) is the linear predictor, and $\phi_{SiteStatus}$ is the precision parameter (dispersion) specific to the level of Site Status.

**Supporting information S2: Temporal Models notation**

In all models below $\mu_i$ is the predicted mean, $u_j$ is the random intercept for Site ID j and $\tau^2$ is the random effect variance. In all model the random effect variance was assumed to be normally distributed with: $u_j \sim N(0, \tau^2)$

**1) Peat Depth over Time**

This model uses a linear fixed effect for year. And peat depth was assumed to be normally distributed with: $\text{Peat Depth}_i \sim N(\mu_i, \sigma^2)$

$$mu_i = \beta_0 + \beta_1 \cdot \text{year}_i + u_{j[i]}$$

**2) Species Richness over Time**

This model used a smooth, unconstrained function for year, s(year), with $\text{Species Richness}_i \sim \text{Poisson}(\lambda_i)$ and

$$log(\lambda_i) = \beta_0 + s(\text{year}_i) + u_{j[i]}$$

**3) Specialist Richness over Time**

This model used a smooth, unconstrained function for year, s(year), constrained to k=5 basis functions. Specialist richness was assumed to be Poisson distributed: $\text{Specialist Richness}_i \sim \text{Poisson}(\lambda_i)$ and:

$$log(\lambda_i) = \beta_0 + s(\text{year}_i, k = 5) + u_{j[i]}$$

**4) Specialist Cover over Time**

This model uses a smooth function for year that is monotonically increasing (mpi) and uses a logit link for the quasibinomial distribution: $\text{Specialist\_cover\_proportion}_i \sim \text{Quasibinomial}(\mu_i)$

$$\text{logit}(\mu_i) = \beta_0 + s_{\text{mpi}}(\text{year}_i, k = 5) + u_{j[i]}$$

### Model diagnostic plots

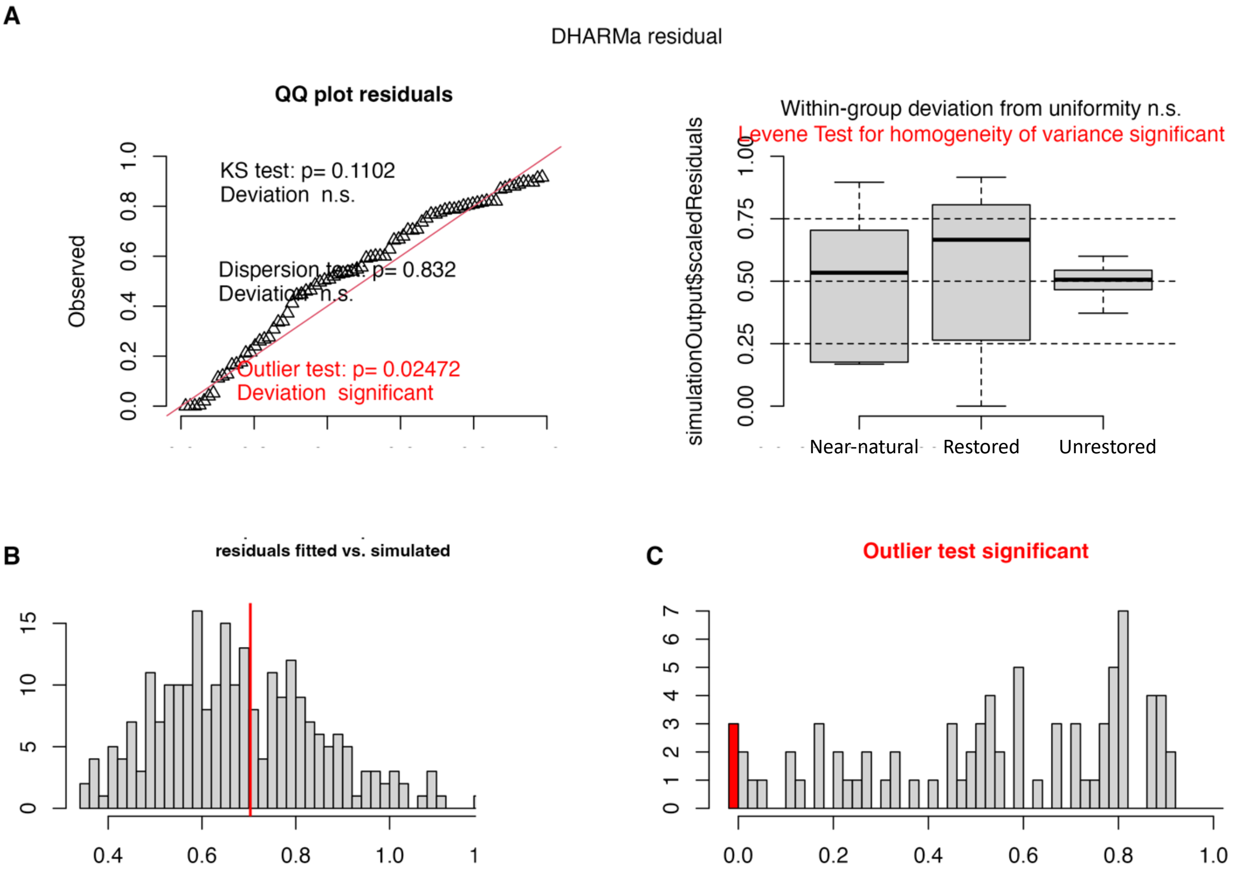


**Figure S2**: Model diagnostics for the model investigating the overall differences in peat depth between unrestored, restored and semi-natural small fens (Gaussian, log1p scale, heteroscedastic residuals by site type: unrestored, restored, near-natural). These plots assess model fit, residual distribution, variance homogeneity among site types, and outlier presence using the simulated residuals within the DHARMa package (Hartig 2025). (**A**) Left: QQ plot of scaled residuals with Kolmogorov-Smirnov (KS) and dispersion tests (both n.s.). Right: The boxplot shows within-group deviation from uniformity (n.s.) and a significant Levene test for heterogeneity of variance (red), grouped by site type – even after accounting for heteroscedastic residuals. This is mainly since peat depth on unrestored sites was usually minimal. Thus, this should not affect the conclusions from the results (**B**) Histogram of residuals (fitted vs. simulated). (**C**) Histogram of residuals with significant outlier test (red).

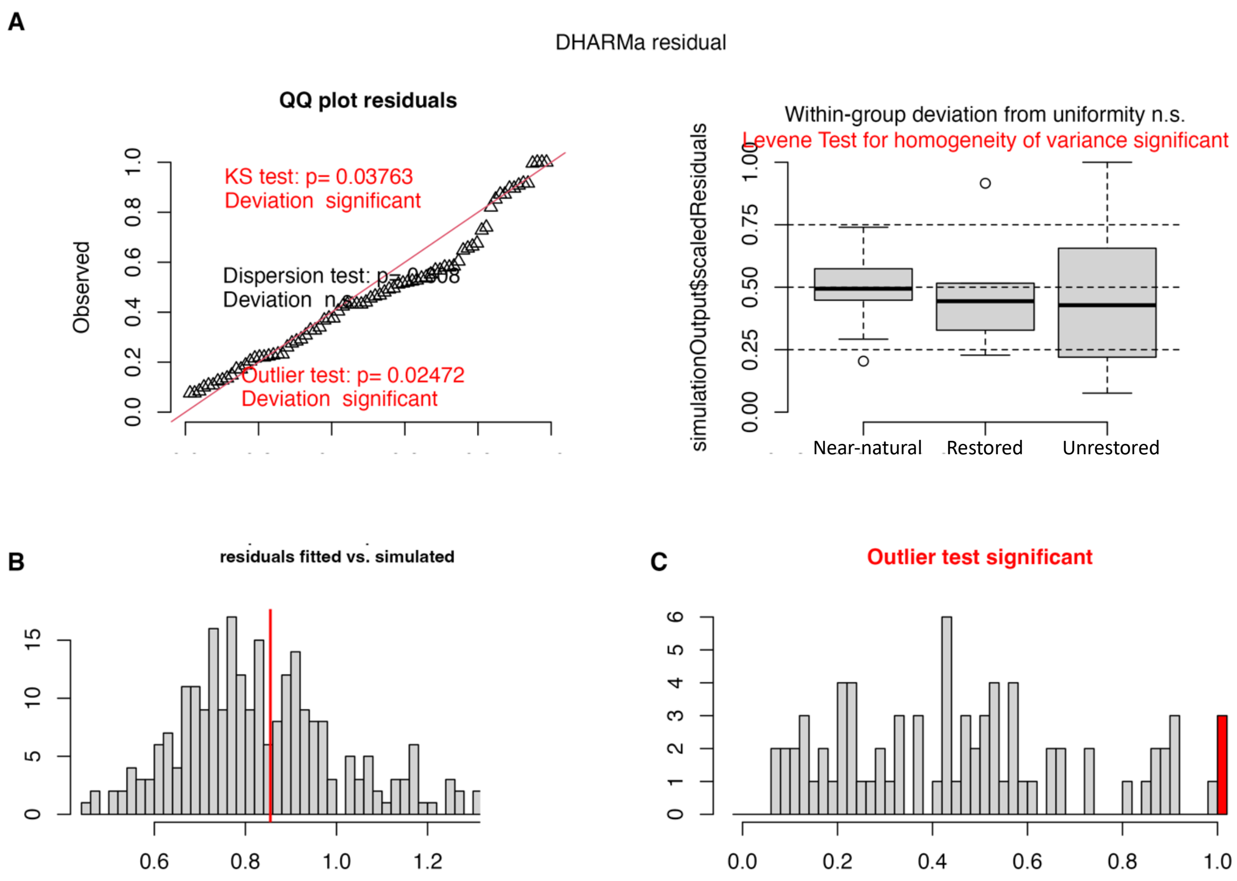


**Figure S3**: Model diagnostics for the model investigating the overall differences in species richness between unrestored, restored and semi-natural small fens (Poisson GLMM on transformed scale; homoscedastic residuals by site type: near-natural, restored, unrestored). These plots assess model fit, residual distribution, variance homogeneity among site types, and outlier presence using the simulated residuals within the DHARMa package (Hartig 2025). (**A**) Left: QQ plot of scaled residuals with Kolmogorov-Smirnov (KS test p = 0.038), and dispersion tests (both n.s.), but a significant outlier test (p=0.02472, red). Right: The boxplot shows within-group deviation from uniformity (n.s.) and a significant Levene test for heterogeneity of variance (red), grouped by site type. (**B**) Histogram of residuals (fitted vs. simulated). (**C**) Histogram of residuals with significant outlier test (red).

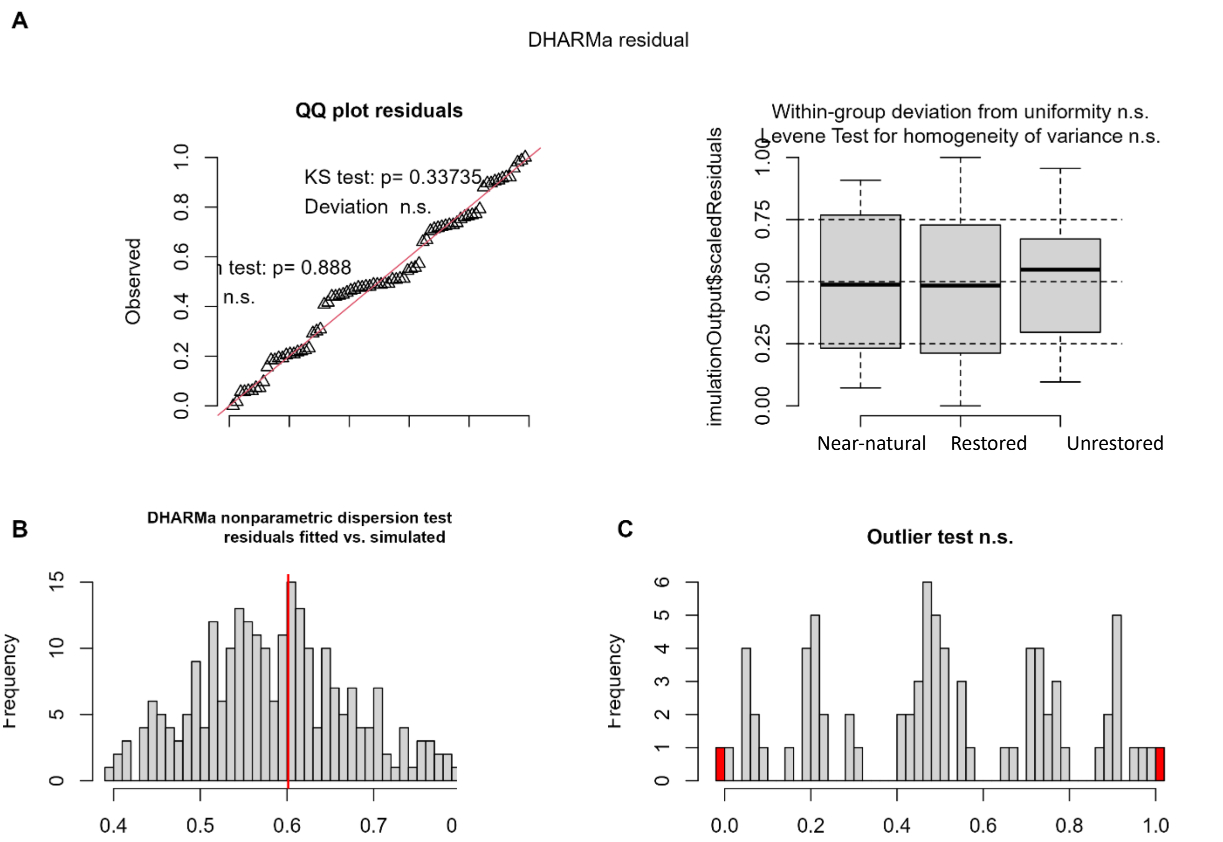


**Figure S4**: Model diagnostics for the model investigating the overall differences in fen specialist species richness between unrestored, restored and semi-natural small fens (Poisson GLMM on transformed scale; homoscedastic residuals by site type: near-natural, restored, unrestored). These plots assess model fit, residual distribution, variance homogeneity among site types, and outlier presence using the simulated residuals within the DHARMa package (Hartig 2025). (**A**) Left: QQ plot indicates no significant deviation from uniformity and dispersion test. Right: The boxplot shows no within-group deviation from uniformity and no heterogeneity of variance between site types. (**B**) Histogram of residuals (fitted vs. simulated). (**C**) Histogram of residuals with significant outlier test (red).

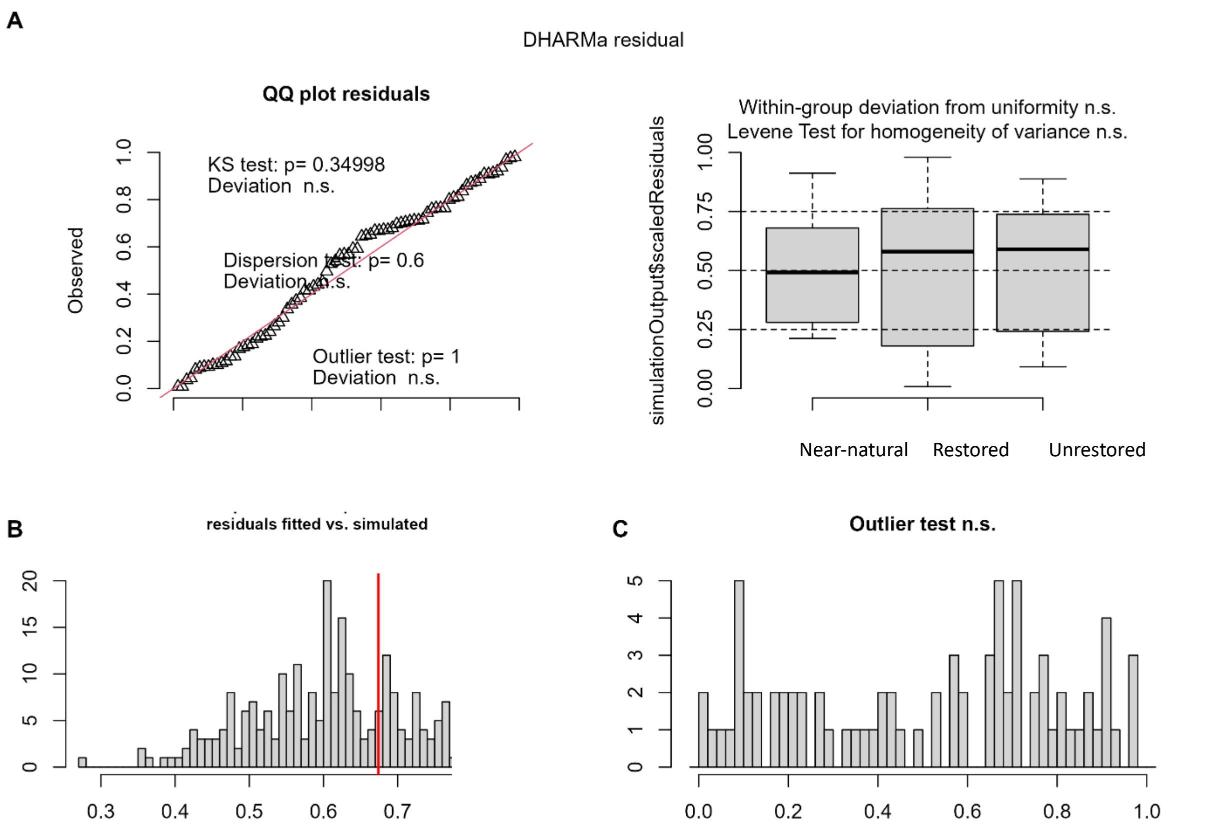


**Figure S5**: Model diagnostics for the model investigating the overall differences in fen specialist species cover between unrestored, restored and semi-natural small fens (Beta GLMM on transformed scale; heteroscedastic residuals by site type: near-natural, restored, unrestored). These plots assess model fit, residual distribution, variance homogeneity among site types, and outlier presence using the simulated residuals within the DHARMa package (Hartig 2025). (**A**) Left: QQ plot indicates no significant deviation from uniformity. Right: The boxplot shows no within-group deviation from uniformity and no heterogeneity of variance between site types. (**B**) Histogram of residuals (fitted vs. simulated). (**C**) Histogram of residuals with significant outlier test (red).

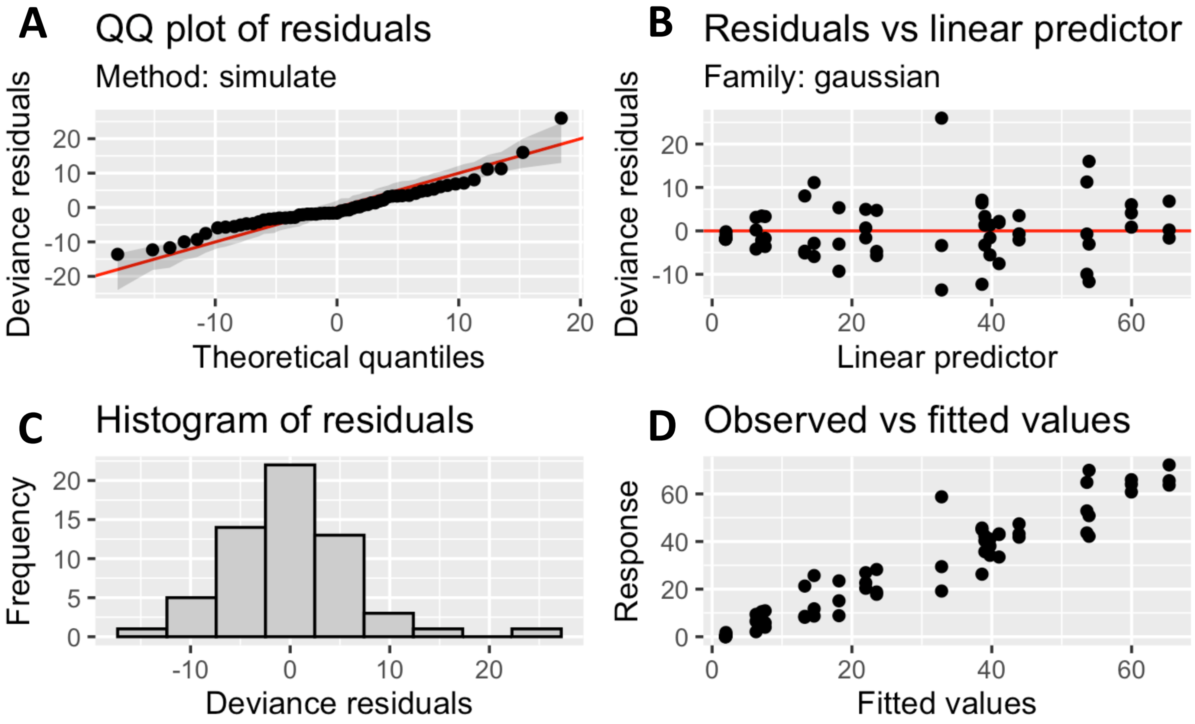


**Figure S6**: Model diagnostic plots of the generative additive model (GAM) investigating the effect of restoration on peat depth over time. Produced with the draw() function in the *gratia* package (Simpson 2024). Model diagnostic plots showing residual distribution and fit. Panels display: **A**) QQ plot of residuals, **B)** residuals versus linear predictor, **C**) histogram of residuals, and **D)** observed versus fitted values for the Gaussian GAM investigating how peat depth changes with time since restoration.

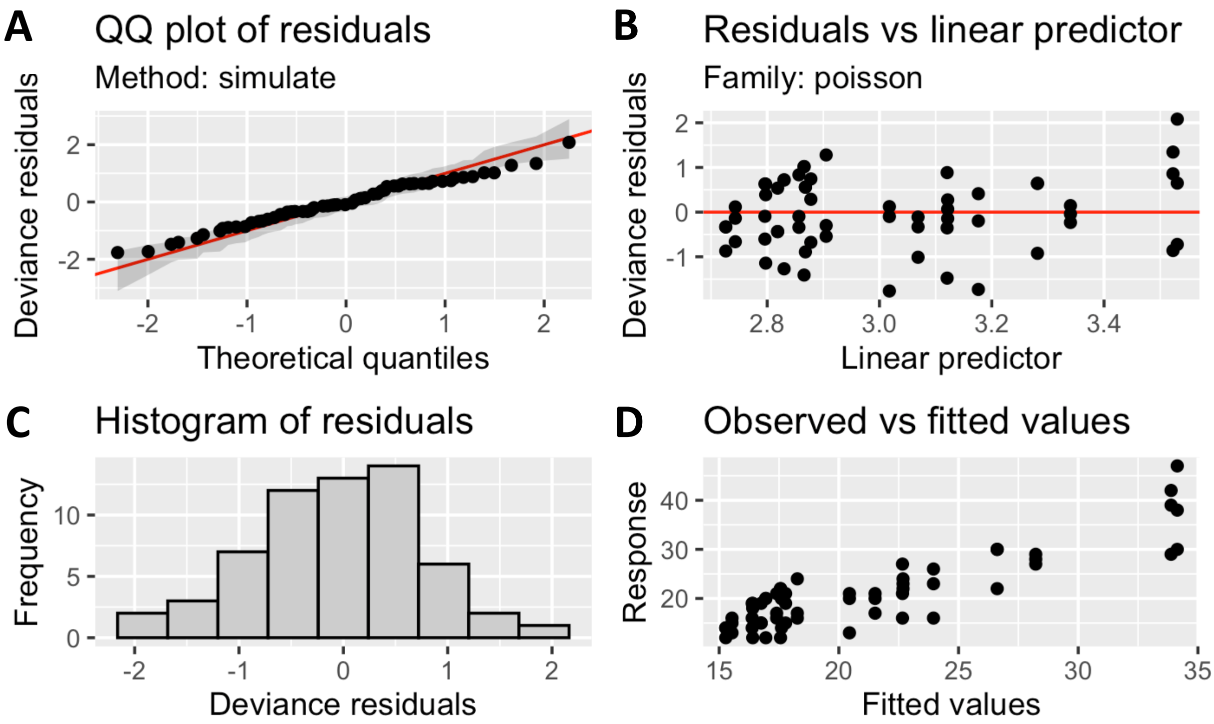


**Figure S7**: Model diagnostic plots of the generative additive model (GAM) investigating the effect of restoration on plant pecies richness over time. Produced with the draw() function in the *gratia* package (Simpson 2024). Model diagnostic plots showing residual distribution and fit. Panels display: **A**) QQ plot of residuals, **B)** residuals versus linear predictor, **C**) histogram of residuals, and **D)** observed versus fitted values for the Gaussian GAM investigating how peat depth changes with time since restoration.

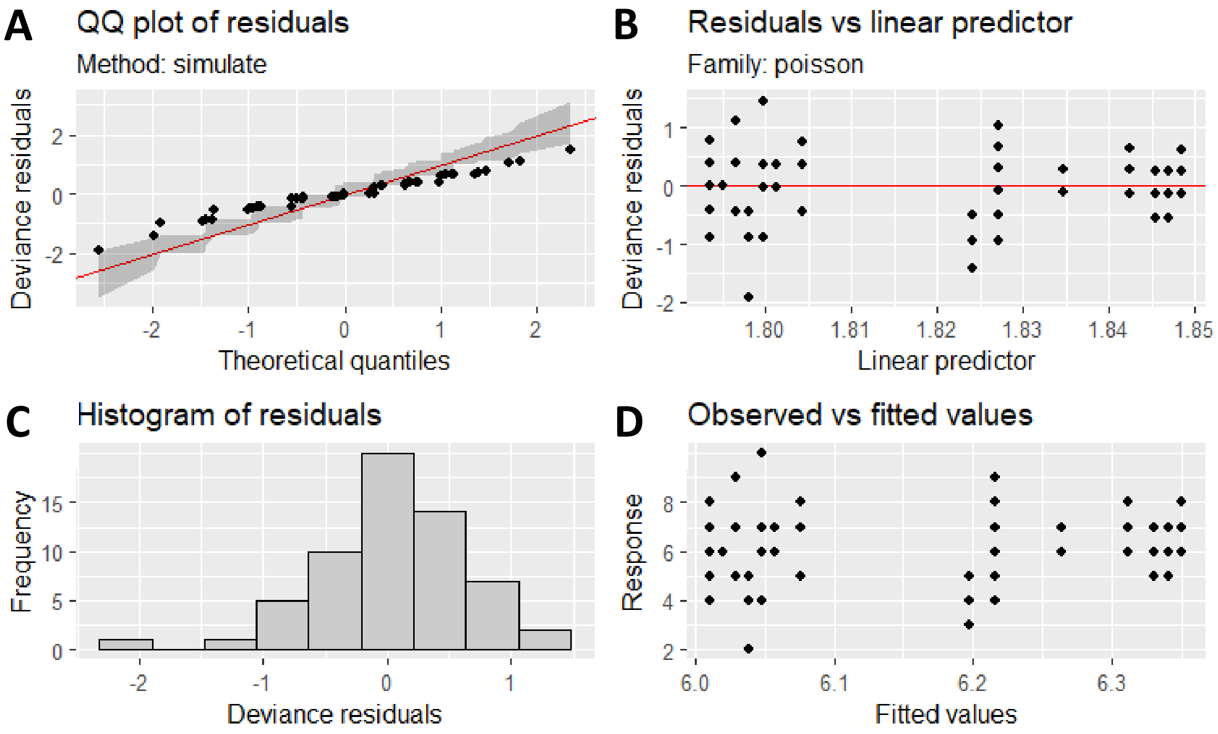


**Figure S8**: Model diagnostic plots of the generative additive models (GAM) investigating the effect of restoration on fen specialist species richness over time. Produced with the draw() function in the *gratia* package (Simpson 2024). Model diagnostic plots showing residual distribution and fit. Panels display: **A**) QQ plot of residuals, **B)** residuals versus linear predictor, **C**) histogram of residuals, and **D)** observed versus fitted values for the Gaussian GAM investigating how peat depth changes with time since restoration.

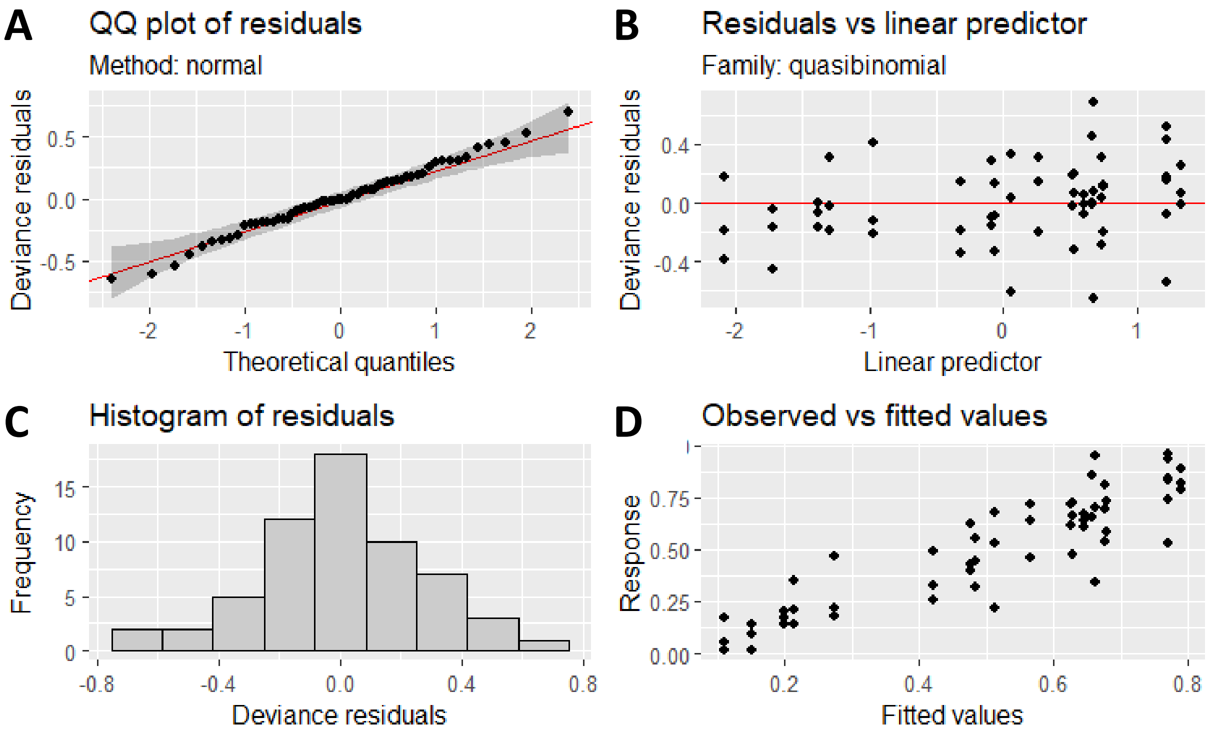


**Figure S9**: Model diagnostic plots of the generative additive model (SCAM) investigating the effect of restoration on fen specialist species cover over time. Produced with the draw() function in the *gratia* package (Simpson 2024). Model diagnostic plots showing residual distribution and fit. Panels display: **A**) QQ plot of residuals, **B)** residuals versus linear predictor, **C**) histogram of residuals, and **D)** observed versus fitted values for the Gaussian GAM investigating how peat depth changes with time since restoration.

**Temporal Models summary**

**Table S2**: Summary of Fixed-Effect Results from Generalized Additive Models (GAMs) and Shape-Constrained Additive Models (SCAM) assessing the effect of year on ecological variables. We estimated "fixed-only" $R^2$ by refitting each model without random-effect smooths and extracting the adjusted $R^2$ from the refit. "Total" fit for GAMs was taken as the adjusted $R^2$ from the original model including random-effect smooths.

| Response variable | Effect | Test Statistic | Test value | p-value | family | link | $R^2$ (fixed) | $R^2$ (total) |
|---|---|---|---|---|---|---|---|---|
| Peat depth | year | t | 6.084 | <0.001 | gaussian | identity | 0.61 | 0.87 |
| Species richness | s(year) | Chi.sq | 27.701 | <0.001 | poisson | log | 0.48 | 0.68 |
| Fen specialists | s(year) | Chi.sq | 0.15 | 0.696 | poisson | log | -0.004 | -0.004 |
| Specialist cover | s(year) | F | 18.37 | <0.001 | quasibinomial | logit | 0.41 | 0.73 |
